# Serial Crystallography with Multi-stage Merging of 1000s of Images

**DOI:** 10.1101/141770

**Authors:** Alexei S Soares, Yusuke Yamada, Jean Jakoncic, Sean McSweeney, Robert M Sweet, John Skinner, James Foadi, Martin R. Fuchs, Dieter K. Schneider, Wuxian Shi, Babak Andi, Lawrence C Andrews, Herbert J Bernstein

## Abstract

KAMO and Blend provide particularly effective tools to manage automatically the merging of large numbers of datasets from serial crystallography. The requirement for manual intervention in the process can be reduced by extending Blend to support additional clustering options such as use of more accurate cell distance metrics and use of reflection-intensity correlation coefficients to infer “distances” among sets of reflections. This increases the sensitivity to differences in unit cell parameters and allows for clustering to assemble nearly complete datasets on the basis of intensity or amplitude differences. If datasets are already sufficiently complete to permit it, one applies KAMO once and clusters the data using intensities only. If starting from incomplete datasets, one applies KAMO twice, first using cell parameters. In this step we use either the simple cell vector distance of the original Blend, or we use the more sensitive NCDist. This step tends to find clusters of sufficient size so that, when merged, each cluster is sufficiently complete to allow reflection intensities or amplitudes to be compared. One then uses KAMO again using the correlation between the reflections having a common hkl to merge clusters in a way sensitive to structural differences that may not have perturbed the cell parameters sufficiently to make meaningful clusters.

Many groups have developed effective clustering algorithms that use a measurable physical parameter from each diffraction still or wedge to cluster the data into categories which then can be merged, one hopes, to yield the electron density from a single protein form. Since these physical parameters are often largely independent from one another, it should be possible to greatly improve the efficacy of data clustering software by using a multi-stage partitioning strategy. Here, we have demonstrated one possible approach to multi-stage data clustering. Our strategy is to use unit-cell clustering until merged data is sufficiently complete then to use intensity-based clustering. We have demonstrated that, using this strategy, we are able to accurately cluster datasets from crystals that have subtle differences.

## 1. Introduction

KAMO (Yamashita, 2017) (Yamashita *et al*., 2017) (Hasegawa *et al*., 2017) (Yamashita *et al*., 2018) and Blend (Foadi *et al*., 2013) provide particularly effective tools to automatically manage the merging of large numbers of datasets from serial crystallography. The requirement for manual intervention in the process can be reduced by extending Blend to support additional clustering options thereby increasing the sensitivity to differences in unit cell parameters and allowing for clustering to assemble nearly complete datasets on the basis of intensity or amplitude differences. KAMO provides the necessary process-flow management infrastructure. This process flow is shown in Fig. 1. If datasets are already sufficiently complete to permit it, apply KAMO once and cluster the data using intensities only. If starting from incomplete datasets, one applies KAMO twice, first using cell parameters. In this step either the simple cell vector distance of the original Blend is used, or the more sensitive NCDist, to find clusters to merge to achieve sufficient completeness to allow intensities or amplitudes to be compared. One then uses KAMO again using the correlation between the reflections having common hkls (Assmann *et al*., 2016) to merge clusters in a way sensitive to structural differences that may not have perturbed the cell parameters sufficiently to make meaningful clusters.

**Fig. 1.**
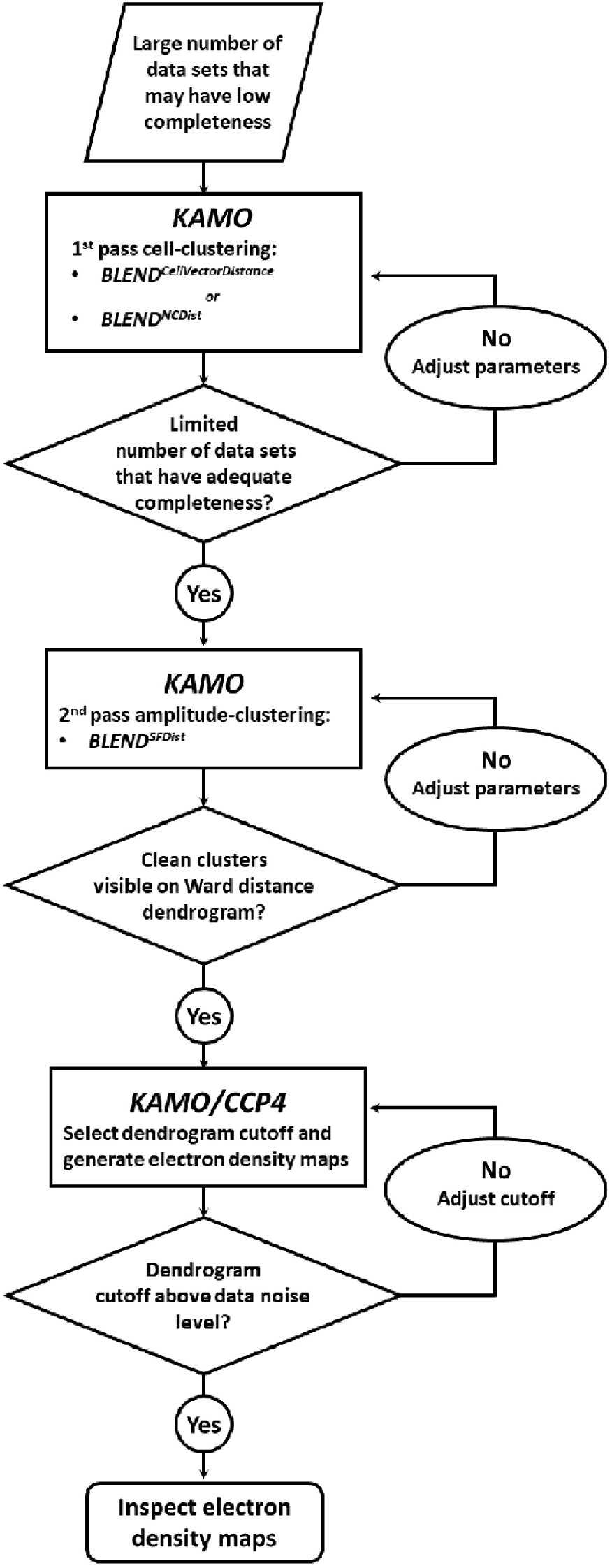
Process flow in use of KAMO and Blend. In the case of the four-way clustering discussed in §3 and §4, a total of 896 data sets were input to the first stage NCDist clustering engine and a total of 107 data sets were input to the second stage SFDist clustering engine (first and second rectangles in Fig. 1).

X- ray free-electron lasers (XFELs) have pioneered effective crystallographic data collection from large numbers of crystals (Colella & Luccio, 1984) (Neutze *et al*., 2000). Serial crystallography, an essential technique at x-ray free electron laser light sources, has become an important technique at synchrotrons (Giordano *et al*., 2012) (Liu & Hendrickson, 2013) (Rossmann, 2014) (Standfuss & Spence, 2017), especially at newer high-intensity synchrotron beamlines (Pearson & Mehrabi, 2020). The data may be organized either as XFEL-like still images or as thousands of wedges of data produced from very large numbers of crystals. The stills and wedges then need to be carefully organized into reasonably homogeneous clusters of data that can be merged for processing. This is going to be one of the common tools to assemble complete data from many partial wedges in MR, SAD, and ligand studies and to sort classes of crystals for studies of dynamics, binding, interactions, *etc.* KAMO includes cluster analysis based both on cell parameters and on reflection correlation coefficients. The clustering is based on Ward’s Method, “Ward’s minimum variance criterion minimizes the total within-cluster variance. To implement this method, at each step find the pair of clus-ters that leads to minimum increase in total within-cluster variance after merging. This increase is a weighted squared distance between cluster centers.” (Wikipedia, 2022)

In this paper we discuss the issues involved in improving the sensitivity of both approaches to clustering, using, as an example, 999 5*^◦^* wedges from lysozyme in four forms:

- NAG: native with N-acetylglucosamine (NAG) soaked in,
- benzamidine: native with benzamidine soaked in,
- benzamidine plus NAG: native with both NAG and benzamidine soaked in.
- Native: no ligands

As we will see, although the cell parameters are changed sufficiently to allow recognition of the NAG soak, it is difficult to filter the benzamidine soak simply on the basis of cell parameter changes, suggesting the desirability of switching from cell-based clustering to intensity-based clustering as early in the process as possible. Hence, cell-based clustering is more universal, in the sense that we can successfully apply it very early in the structure solution process, but it is less granular in the sense that it cannot see as much detail as intensity-based clustering. Cell-based clusters are less able to discriminate between similar but non-identical forms.

Two-stage clustering can be regarded as a method to reduce the data multiplicity needed to achieve a desired level of data precision. Hence, it is a continuation of previous techniques (White *et al*., 2012) that reduced the need for multiplicity compared to the Monte Carlo method of integration (Kirian *et al*., 2010), which makes no assumptions regarding crystal-to-crystal scaling (and hence relies entirely on statistical averaging to achieve data precision from millions of observations).

An alternative to consider, rather than staging the cell-based clustering first and then applying intensity-based clustering, would be to combine the two approaches in a single stage of higher dimensionality. There are two problems that would have to be addressed in order to create such a single-stage combined algorithm.

First, it is not possible to apply intensity-based clustering at all without first indexing all reflections in a way that provides a common unambiguous label for each reflection in each image to correctly identify corresponding reflections with the same index in each different image. One option to satisfy this requirement is to index all diffraction images in P1 or some other low symmetry common to all images. This may require more images and more reflections than are available and, worse, because symmetry is being ignored, may bring together for merging images describing very different molecular conformations.

Second, as the number of independent parameters rises, all clustering methods work increasingly poorly due to the “curse of dimensionality” (Bellman, 1956). Combining the very-well-behaved low-dimensionality cell-based clustering with the high-dimensionality intensity-based clustering can obscure the results of cell-based clustering. In most cases, unless the information to be gained from cell-based clustering is available *a priori*, it is more useful to process the cell-based clustering information first and then to move on to intensity-based clustering in a second stage based on the cell-based results.

For cases in which the indexing is too ambiguous to reliably start the process, the approach used in dials.cosym should be considered (Gildea & Winter, 2018), which uses the averages of intensities within images and from multiple images simultaneously with spot positions for indexing, dealing with the issue of the curse of dimensionality by using the averaging to limit the increase in dimension as much as possible (Brehm & Diederichs, 2014) (Diederichs, 2017). This is not the same as simultaneously doing full cell-based and full intensity-based clustering, which is not recommended.

## 2. Limits of conventional clustering

Since our goal was to expand the capabilities of existing clustering techniques, we began by applying a conventional clustering strategy to diffraction data from lysozyme micro-crystals containing various combinations of known small molecule binders. Micro- crystals were preferred to avoid the conflation of structurally anisotropic data that has been demonstrated in larger crystals; see (Thompson *et al*., 2018).

Lysozyme micro-crystals suitable for acoustic harvesting (Soares *et al*., 2011) were grown using batch crystallization by dissolving 120 mg/ml lysozyme in 0.2M sodium acetate pH 4.6 (Hampton Research HR7-110) and combining with equal parts precipitant (10% ethylene glycol + 12% sodium chloride) (Roessler *et al*., 2016). The resulting slurry of 5 – 10 *µ*m crystals was divided into four aliquots. Three of the four aliquots were then equilibrated overnight with an equal volume of 0.5 M solutions of, respectively, benzamidine, NAG, and benzamidine plus NAG. These two small molecules are known to bind tetragonal lysozyme crystals (Yin *et al*., 2014). The fourth aliquot was diluted with an equal volume of mother liquor but contained no ligands.

The diffusion rate for benzamidine and NAG within lysozyme crystals is approximately 1*µm/s* (Cole *et al*., 2014). To prevent cross-contamination of crystals with neighboring forms, crystals could not be mixed with different forms for more than 1s before diffusion was halted by plunge cryo-cooling in *LN*_2_. To accomplish this, we deposited 5 *µL* of crystal slurry from each aliquot onto a separate agarose support (Cuttitta *et al*., 2015). We used acoustic sound pulses to harvest 2.5 *nL* of crystal slurry from each of the four lysozyme aliquots and separately positioned them on a micro-mesh (MiTeGen M3-L18SP-10) so that none of the droplets was in contact with any other. Crystal-containing droplets were threaded through small apertures to prevent cross-contamination (Foley *et al*., 2016). We then swept the non-crystal-containing side of the micro-mesh against a sponge moistened with cryo-protectant (mother liquor + 20% glycerol) and, in one smooth motion, immediately cryo-cooled the micro-mesh in *LN*_2_. In addition to cryo-protection, this also mixed the crystals together into one contiguous field. The same procedure was repeated for a micro-mesh containing only two lysozyme forms, benzamidine plus NAG and native. Serial diffraction data were then obtained in 5 degree wedges from approximately 100 crystals on each micro-mesh.

The software package KAMO was then used in default configuration (in which data are clustered based only on unit cell similarity) to partition the diffraction data from micro-meshes containing the four lysozyme forms into four clusters, and the diffraction data from micro-meshes containing two lysozyme forms into two clusters. Each cluster of data was merged and then phased using the known structure of lysozyme. Subsequently the atomic model was refined using refmac (Winn *et al*., 2003), and an omit difference map was examined using coot in the region where the ligands were expected to bind to the protein surface (Emsley & Cowtan, 2004). The omit difference map was contoured at 1.5 *σ* and displayed using Pymol (DeLano, 2002). The omit maps calculated from the four-way clustering data were not observed to match closely any of the four lysozyme forms known to have been acoustically harvested onto the micro-meshes (data not shown). We concluded from this result that the clustering algorithm was not sufficiently sensitive to differentiate these four classes of very similar crystals using only variations in the observed unit cell parameters. However, the omit maps calculated from the two-way clustering data were a good fit to the expected lysozyme forms (Fig. 2). We concluded from this result that the two-ligand form was sufficiently different from the native form that unit-cell-based clustering could be successful. To do the four-way split, we combined the universality advantage of cell-based clustering (§3) with the granularity advantage of intensity-based clustering (§4).

**Fig. 2.**
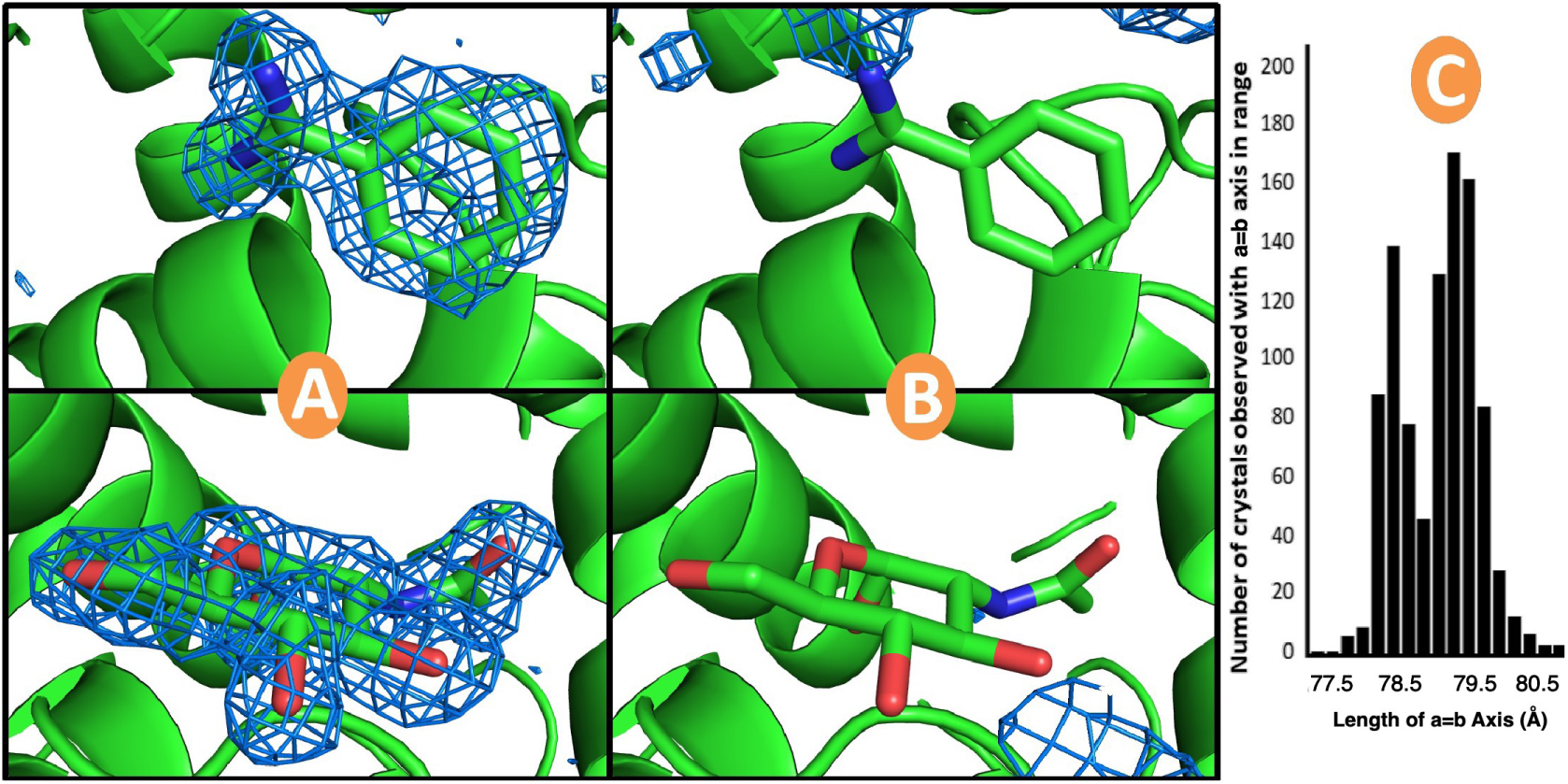
Electron density maps calculated after two-way clustering of diffraction data obtained from micro-meshes that contained a mixture of doubly-bound crystals (benzamidine plus NAG; Fig 2A) and native crystals (no ligands; Fig. 2B). The omit difference maps are contoured at 1.5 *σ* in the region expected to contain benzamidine (top) and NAG (bottom). The histogram cluster in Fig 2C represents the unit cell dimensions of the cluster of crystal datasets that yielded the omit difference map shown in A. Similarly, the histogram cluster on the right in C represents the unit cell dimensions of the cluster of crystal data shown in B. Clearly the clustering algorithm was able to accurately partition the data for this simple two-way split. See A.1 in Supplementary Materials.

## 3. Clustering on Cell Parameters

Stills and wedges of very low completeness are more appropriate for cell parameter clustering, rather than reflection clustering, because pairs of images with very few commensurate reflections may still provide reasonable estimates of unit cells but not provide enough data to compute a meaningful distance between sets of reflections.

The default Blend approach to clustering on cell parameters is to use [*a, b, c, α, β, γ*] as a six-vector, do a principal component analysis (PCA), drop the components without significant variance, and use the Euclidean distance calculated from the remaining components. This approach does not deal as effectively with the discontinuities produced by experimental error and ambiguities in reduction (*e.g.* between Type I and Type II cells and near the cubic unit cells) as does the Andrews-Bernstein Niggli-Cone Distance (NCDist) algorithm (Andrews & Bernstein, 2014). NCDist allows slightly larger clusters of truly similar datasets to be formed, working in the space *G*^6^ which uses Niggli reduction in the six-dimensional space consisting of the metric tensor with the last three components doubled, [*a*^2^*, b*^2^*, c*^2^, 2*bc cos*(*α*), 2*ac cos*(*β*), 2*ab cos*(*γ*)].

In our test case of 999 datasets of lysozyme with NAG and benzamidine soaks 998 clusters are found with completeness ranging from 40% to 100%. The top levels of the two dendrograms are shown in Figs. 3 and 4. The clusters are labelled by Linear Cell Variation (LCV), which “measures the maximum linear increase or decrease of the diagonals on the three independent cell faces. Values below 1% in general indicate a good degree of isomorphism among different crystals. Structural differences start to be noticeable with LCV greater than 1.5%. A value in angstroms associated to LCV is provided by the absolute Linear Cell Variation (aLCV).” (Foadi *et al*., 2013) Note the smaller Ward distances *i.e.* tighter clusters for the equivalent clusters in the latter, NCDist-based dendrogram as compared to the former.

**Fig. 3.**
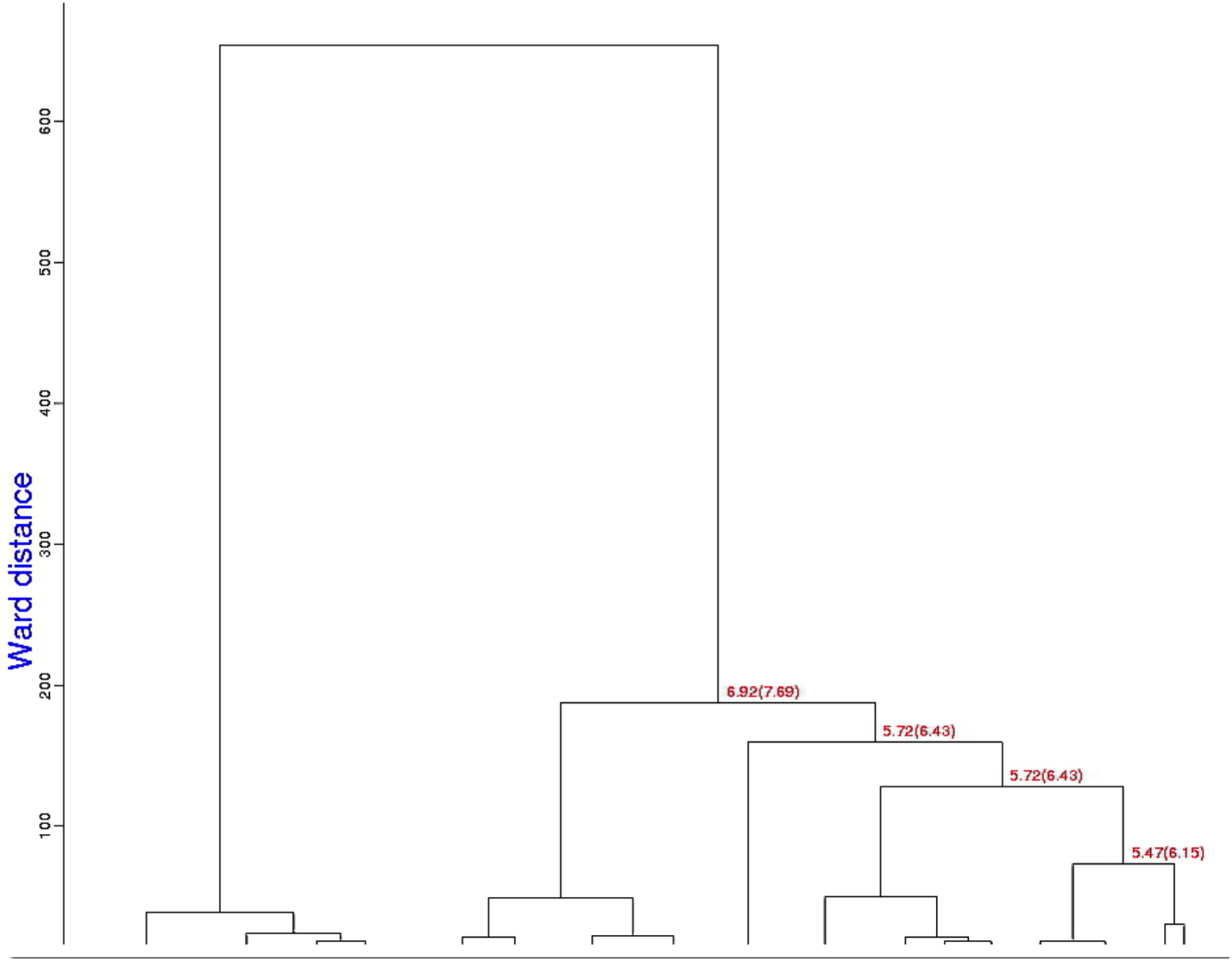
This dendrogram presents the top levels of Blend clustering using the original less-sensitive Blend cell-parameter distance function. The numbers are the LCV and the aLCV, with the aLCV in parentheses.

**Fig. 4.**
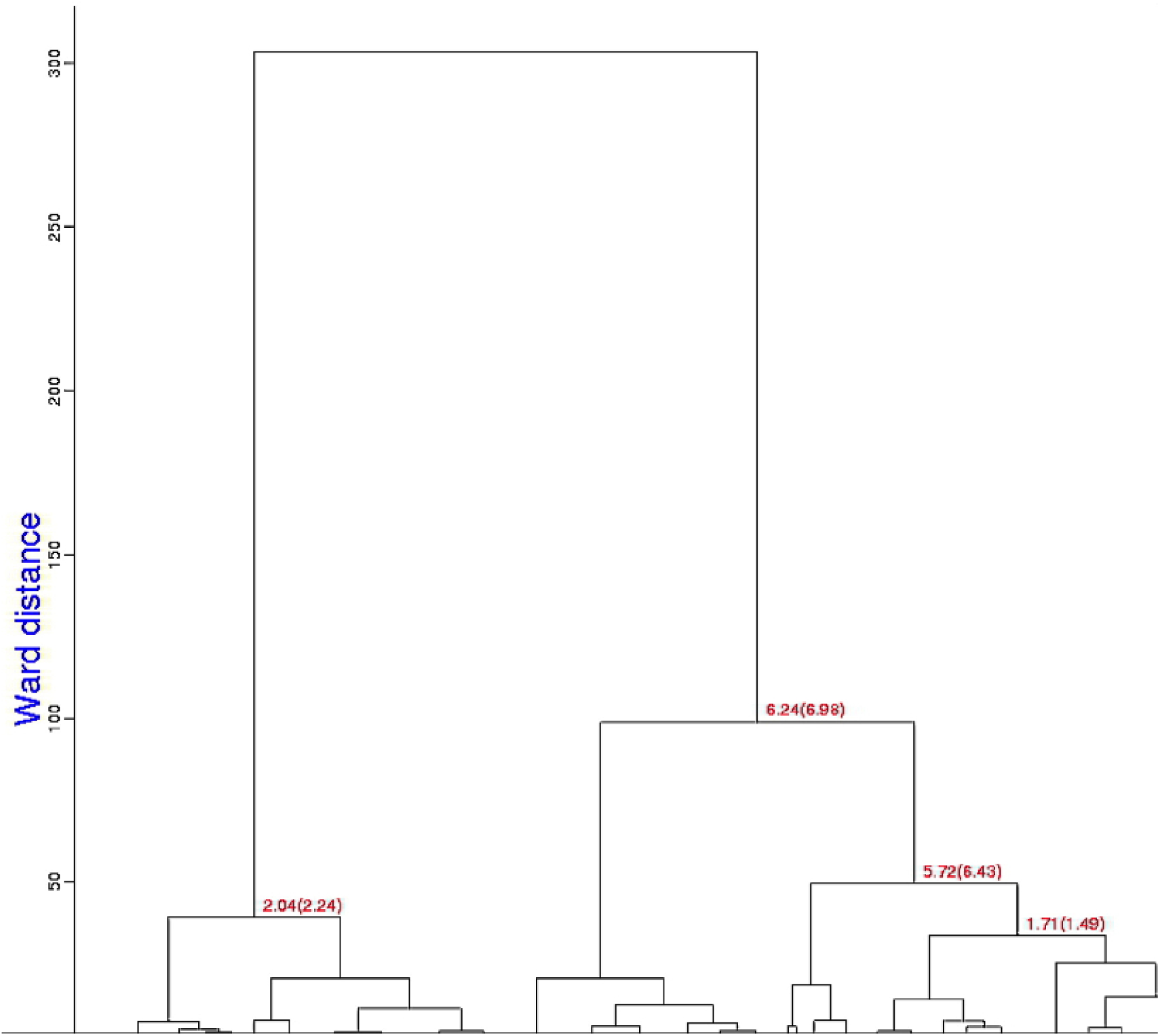
This dendrogram presents the top levels of Blend clustering using the more sensitive Andrews-Bernstein Niggli-Cone-Distance (NCDist) algorithm. The numbers are the LCV and the aLCV, with the aLCV in parentheses. Clustering is guided by the progressive merging of separate clusters into larger ones, using a measure of cluster proximity known as Ward distance. This is equal to the increase of the distance variance (between each element of a cluster and its centroid) resulting from the merging of two separate clusters (Ward, 1963). Note that the Ward distances are smaller than those for the equivalent clusters in Fig. 3.

The dendrograms are qualitatively similar but for this test data the discrimination of the clustering changes. Using the original Blend algorithm, the largest clusters that are 100% native, 100% NAG, 100% benzamidine and 100% benzamidine plus NAG contain 4, 15, 5, and 10 datasets, respectively. Using the NCDist clustering, the largest clusters that are 100% native, 100% NAG, 100% benzamidine and 100% benzamidine plus NAG contain 9, 15, 8, and 7 datasets, respectively. This provides a better base for switching over from cell clustering to reflection clustering; half of the 100% pure clusters are larger using NCDist.

## 4. Clustering on Reflections

In a regime of high completeness, say 90%, different datasets can have enough reflections with common hkl’s to generate a satisfactory similarity or distance for clustering. If the data have been scaled, an R-value can be used as a distance, but for unscaled data the preferred approach is to use a Pearson Correlation Coefficient (*CC*) as a measure of similarity, i.e. having a larger value of the coefficient for sets of reflections that are similar and a smaller value of the coefficient for sets of reflections that are dissimilar. The Pearson Correlation Coefficient is essentially the cosine of the angle between vectors of data. The lack of common scaling is dealt with by subtracting the mean (*µ*) of each vector from each component and dividing by the norm (*||..||*) of each to get two unit length vectors. Recall that the dot product (*·*) of two vectors is equal to the product of the norms of the two vectors times the cosine of the angle between them.

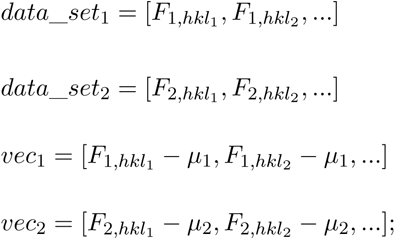

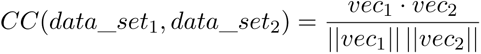

In order to extend the range of applicability of *CC*, we convert it to a distance,

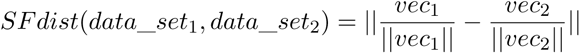

which is related to *CC* by

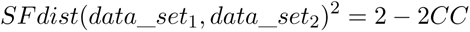

This choice for SFDist allows the distance to be adjusted to account for the greater uncertainty in cases where a pair of data sets has few common reflections, less than 90%, for example, by adding a penalty to the distance for each unmatched reflection.

## 5. Impact of choices in clustering

Unambiguous benzamidine-only, NAG-only, and benzamidine plus NAG clusters are shown in the omit difference maps of the NAG site in clusters 28, 43 and 62 in Figs. 5, 6 and 7, respectively. Omit difference maps of the benzamidine site in clusters 28, 43 and 62 are in Figs. 8, 9 and 10, respectively. These are the results of two-stage KAMO clustering of the test data using NCDist cell-parameter-based clustering to reach 10% completeness and then SFDist reflection-based clustering on the resulting 107 non- overlapping clusters. For an example of this approach when using a higher level of completeness before the cutover from cell-parameter-based clustering to intensity- based clustering, see Nguyen *et al*. (2022).

**Fig. 5.**
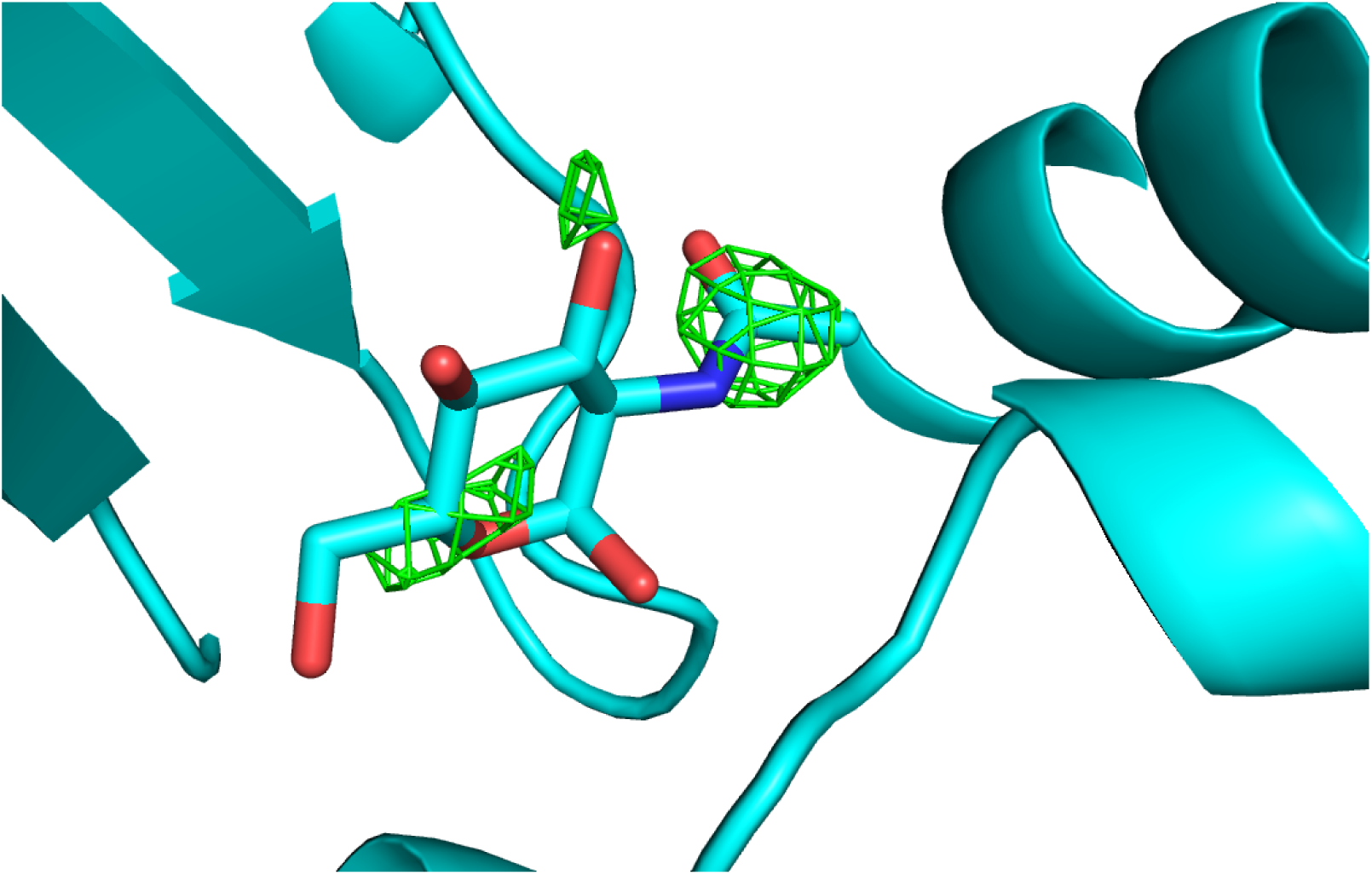
Omit difference map of the NAG site in cluster 28 of a two-stage clustering with KAMO using cell parameters and NCDist to reach 10% completeness and then CC clustering with SFDist.

**Fig. 6.**
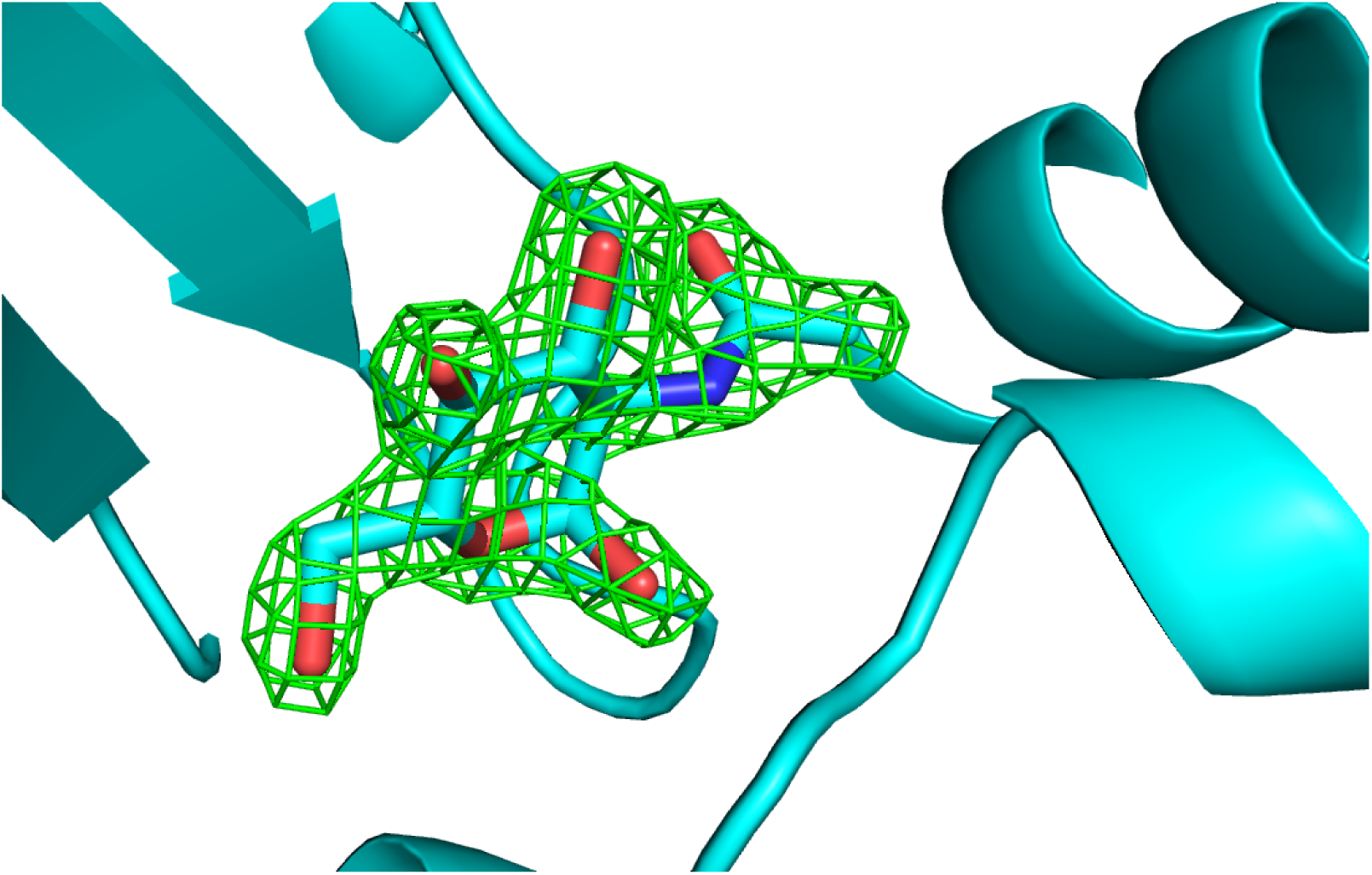
Omit difference map of the NAG site in cluster 43 of a two-stage clustering with KAMO using cell parameters and NCDist to reach 10% completeness and then CC clustering with SFDist.

**Fig. 7.**
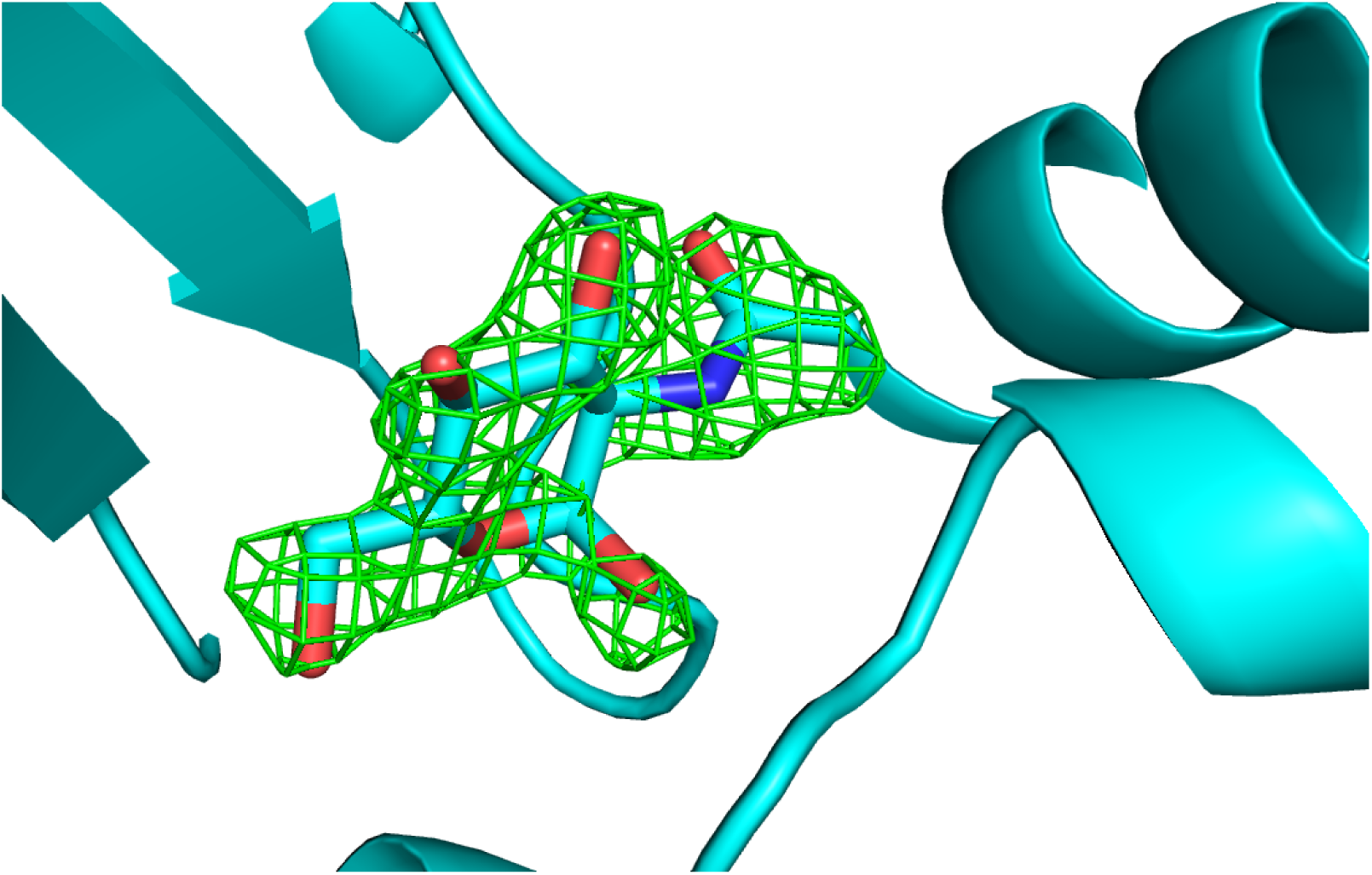
Omit difference map of the NAG site in cluster 62 of a two-stage clustering with KAMO using cell parameters and NCDist to reach 10% completeness and then CC clustering with SFDist.

**Fig. 8.**
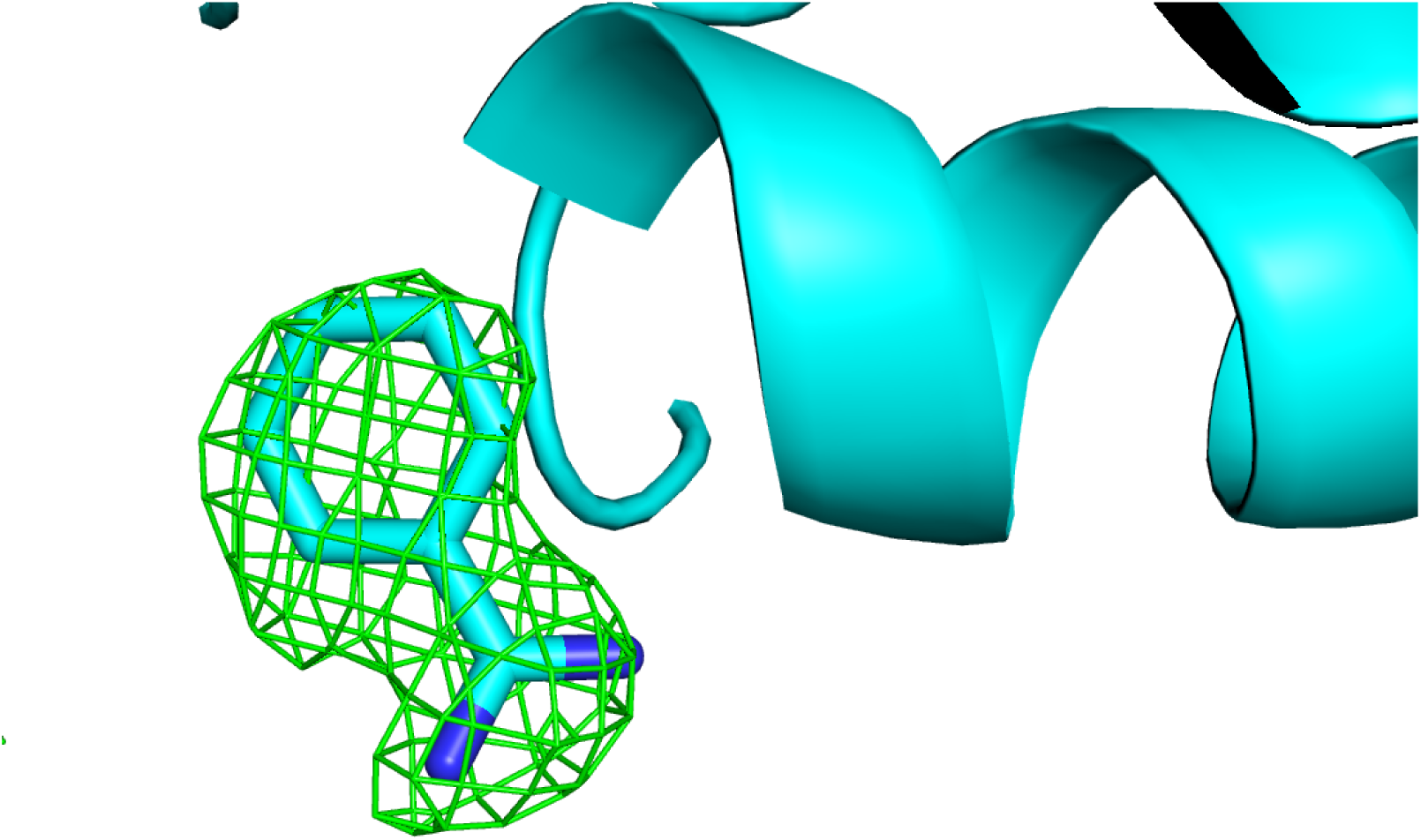
Omit difference map of the benzamidine site in cluster 28 of a two-stage clus- tering with KAMO using cell parameters and NCDist to reach 10% completeness and then CC clustering with SFDist.

**Fig. 9.**
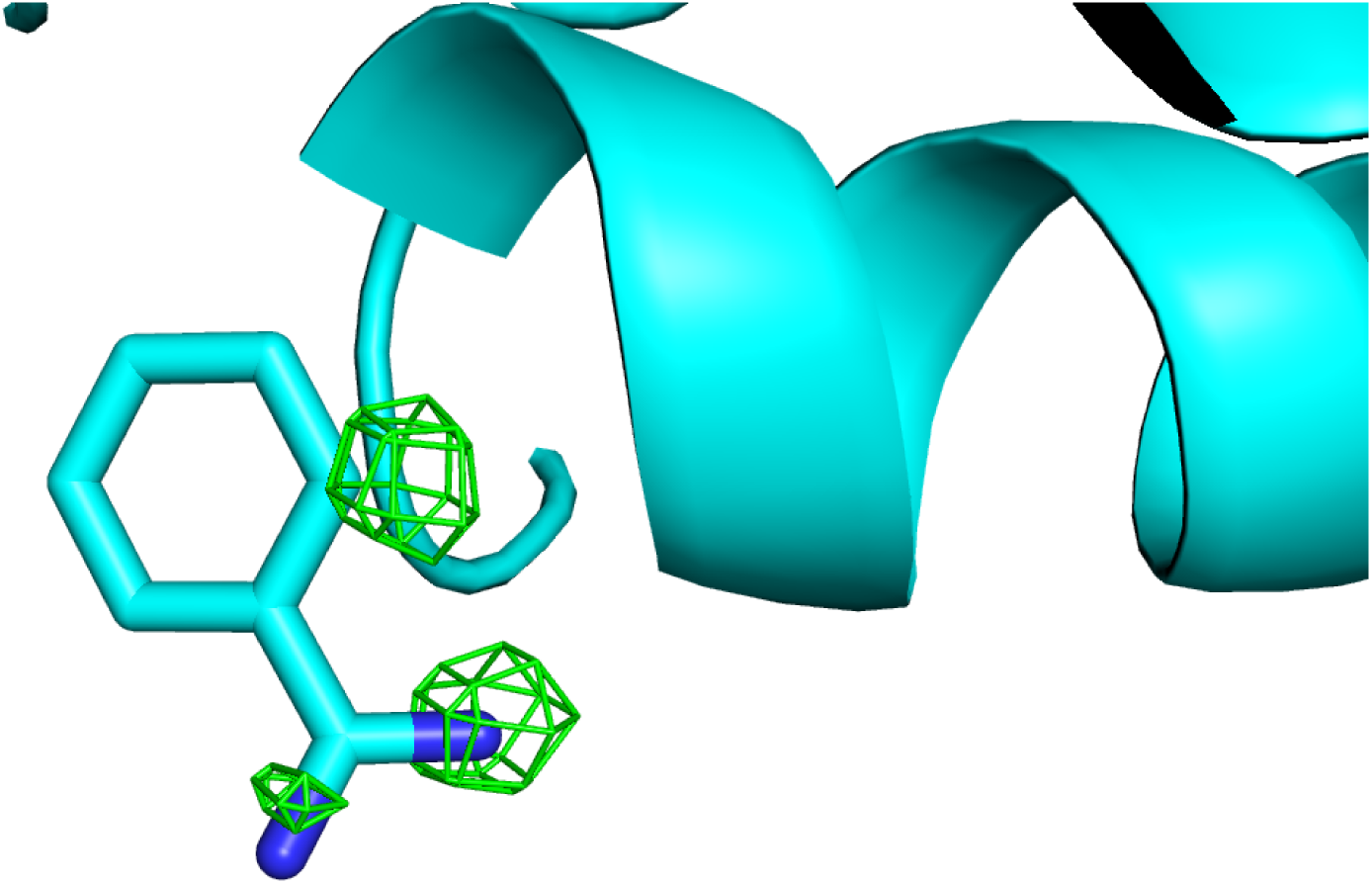
Omit difference map of the benzamidine site in cluster 43 of a two-stage clus- tering with KAMO using cell parameters and NCDist to reach 10% completeness and then CC clustering with SFDist.

**Fig. 10.**
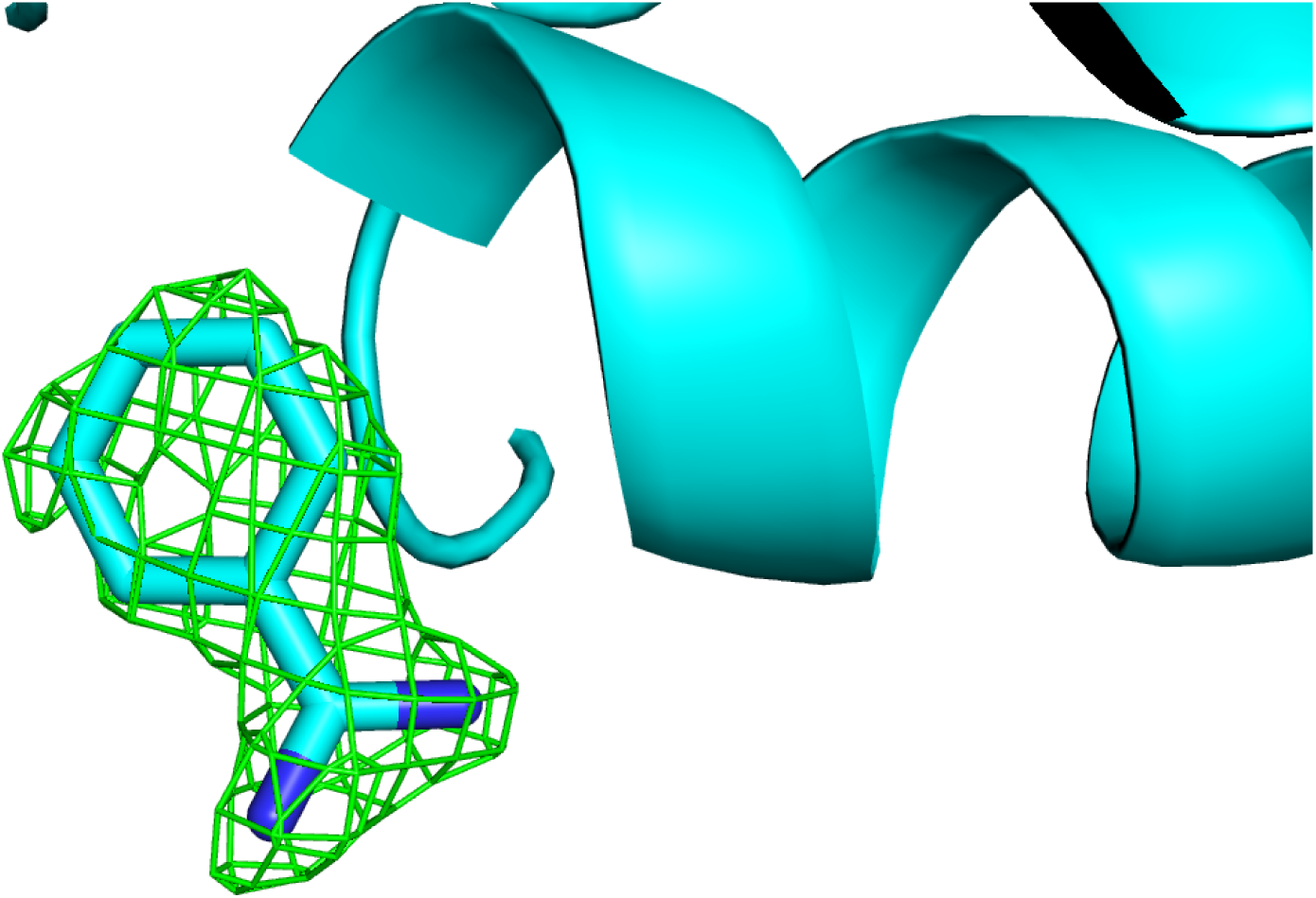
Omit difference map of the benzamidine site in cluster 62 of a two-stage clustering with KAMO using cell parameters and NCDist to reach 10% completeness and then CC clustering with SFDist.

The impact of using clustering on reflections for larger clusters can be seen by looking at how well-represented reasonably pure clusters are. In Figs. 11 and 12, we have represented the purity of native, NAG, benzamidine, and benzamidine plus NAG species using NCDist and SFDist.

**Fig. 11.**
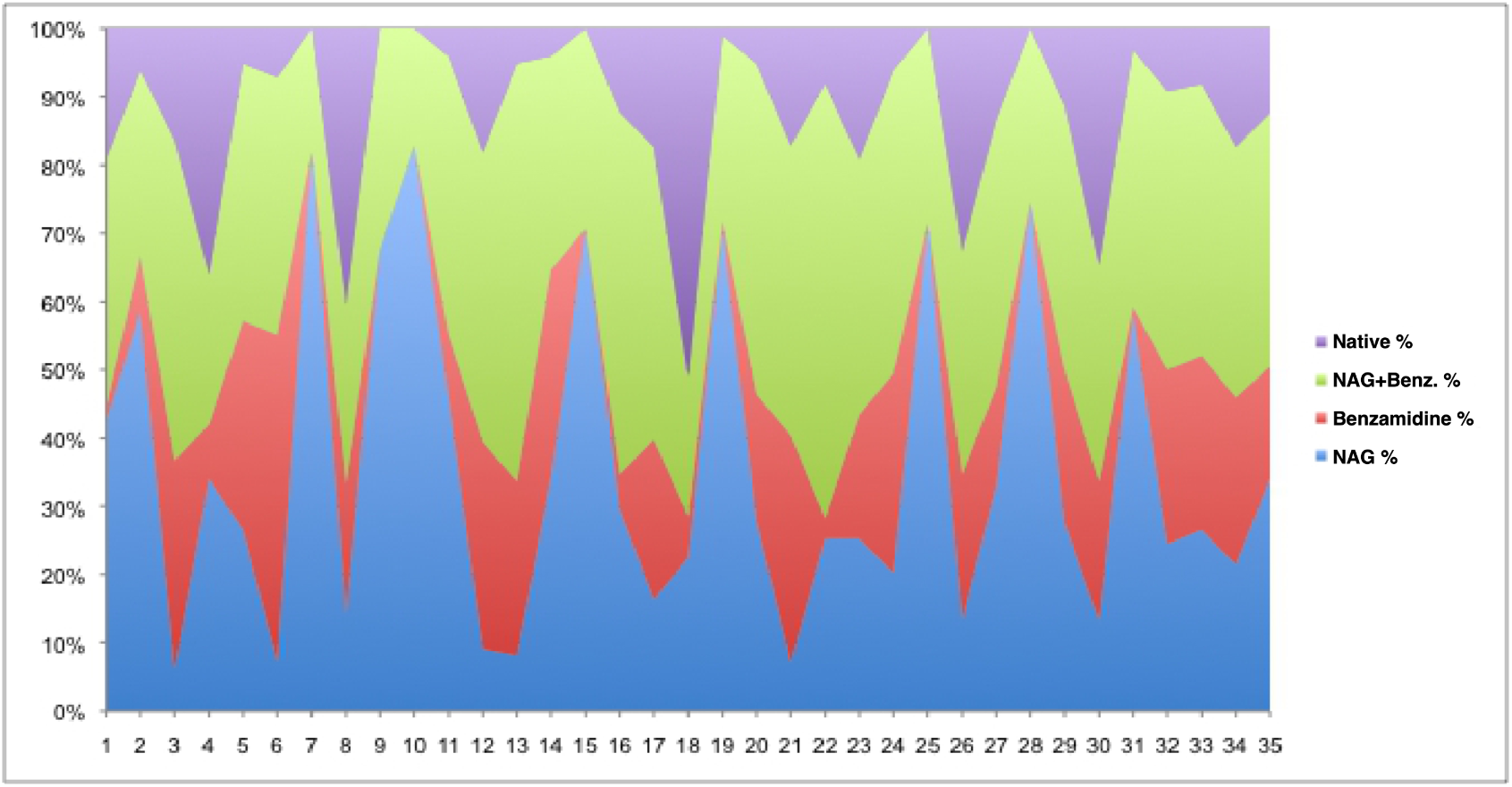
Color charts of the 35 largest dataset clusters for the NCDist clustering. From top to bottom the color blocks are the native soak, the benzamidine plus NAG soak, the benzamidine soak and the NAG soak. If one color reaches nearly from the bottom to the top at a given position, that cluster is a nearly pure species.

**Fig. 12.**
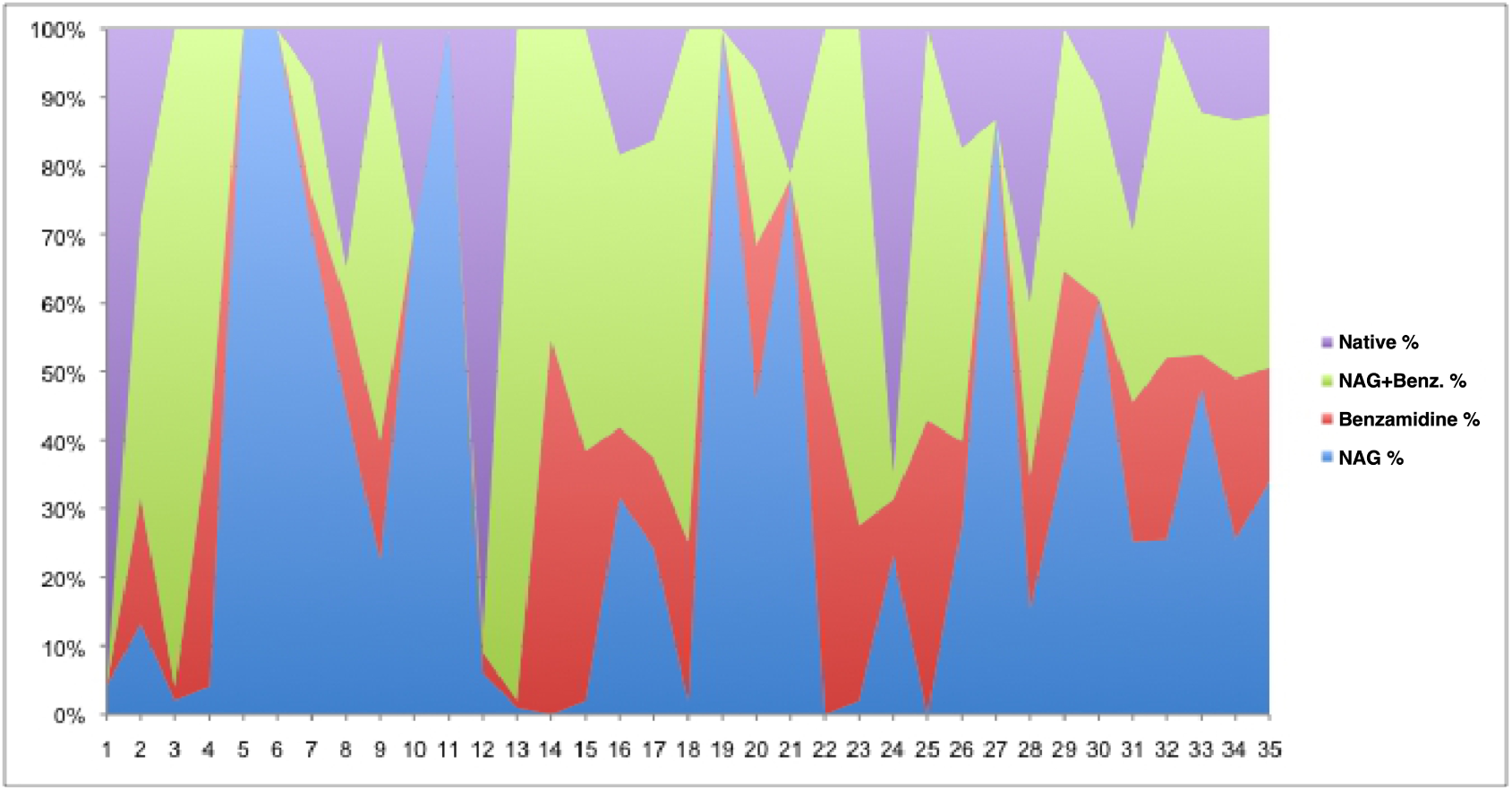
Color charts of the 35 largest dataset clusters for the SFDist clustering. From top to bottom the color blocks are the native soak, the benzamidine plus NAG soak, the benzamidine soak and the NAG soak. If one color reaches nearly from the bottom to the top at a given position, that cluster is a nearly pure species. This is the case for each soak on the left end of this SFDist chart.

The extreme variations in the SFDist results suggest two important lessons.

- It is best to use reflection-based clustering starting from datasets that are small enough to still be likely to be pure species, *i.e.* use cell-based clustering only just far enough to reach sufficient completeness that the intensity-based clustering can be handled.
- It is not necessarily desirable to continue clustering to the largest of the “best” possible clusters. Smaller clusters of sufficient quality for processing are more likely to be pure species.

## 6. Discussion

Because micro-crystals are expected to react quickly and uniformly to changes in their environment, serial crystallography is a desirable tool for examining the plasticity with which protein crystals respond to external perturbations. In some cases the external perturbation can be physical, such as conformational changes induced by light (Young *et al*., 2016). In other cases proteins are perturbed by chemical means (Fromme, 2015). Often it is not possible to to draw a sharp boundary between diffraction images from different protein forms without the assistance of some type of clustering software. In response to this, many groups have developed effective clustering algorithms that use a measurable parameter from each diffraction still or wedge to cluster the data into categories which can then be merged to, hopefully, yield the electron density from a single protein iso-form. Examples of measurable parameters that have been used for this purpose include unit cell dimensions (Foadi *et al*., 2013) (Zeldin *et al*., 2015), and diffraction intensities (Assmann *et al*., 2016) (Diederichs, 2017). What is striking about many of these physical parameters is that they are largely independent from one another.

Consequently, it should be possible to greatly improve the efficacy of data clustering software by combining quasi-independent information in a multi-stage parti-tioning strategy (as presented here). An alternative one might consider in some cases would be to combine the same data in a single higher-dimensional (more independent parameters) stage. However, all clustering methods work increasingly poorly as the number of independent parameters rises due to the “curse of dimensionality” (Bellman, 1956). Combining very well-behaved low-dimensionality cell-based clustering with high-dimensionality intensity-based clustering gives up the advantage of the reliability of cell-based clustering. In most cases, unless all the information to be gained from cell-based clustering is available *a priori*, it is probably best to take advantage of that information first and then move on to intensity-based clustering in a second stage, as we have done here.

We have demonstrated one possible approach to multi-stage data clustering. Our strategy was to use unit-cell-based clustering until merged data was of sufficient completeness to then use intensity-based clustering. We have demonstrated that, using this strategy, we were able to accurately cluster datasets from crystals that had subtle differences. Certainly if one is dealing with a case in which the “correct” symmetry and indexing of all reflections are known for all images, it makes sense to do only intensity-based clustering, but in the general case doing cell-based clustering first make sense.

## 7. Availability of Data

The HKL structure factor files and BLEND clustering data files used for the final intensity-based clustering are available from Zenodo (European Organization For Nuclear Research & OpenAIRE, 2013) (Soares *et al*., 2017). The datasets were collected at National Synchrotron Light Source II (NSLS-II) at the Highly Automated Macromolecular Crystallography (AMX) beamline, 17-ID-1 (Fuchs *et al*., 2016). The coordinates have been deposited at the Protein Data Bank (Bernstein *et al*., 1977) (Berman *et al*., 2000). The datasets are:

- 8DCT Lysozyme from cluster 0003, double apo
- 8DCU Lysozyme cluster 0028, benzamidine ligand
- 8DCV Lysozyme cluster 0043, NAG ligand
- 8DCW Lysozyme cluster 0062, NAG and benzamidine ligands

## Appendix A

### Supplementary Materials

#### A.1. Electron Counting of Ligand Occupancies

The occupancy of benzamidine (BEN) and N-acetylglucosamine (NAG) ligands deduced from the integrated electron density in the volume occupied by each ligand is shown in Fig. A1. The data clusters 3 (Fig. A4), 28 (Fig. A5), 43 (Fig. A6), and 62 (Fig. A7) were obtained using two-stage clustering (BLEND^NCDist^ followed by BLEND^SFDist^), as described in §4. Electron density maps were placed on an absolute scale using iterative density modification (Soares & Caspar, 2017) and the electron density within the envelope of each ligand was integrated. The occupancy was deduced by comparing the observed number of electrons within each ligand envelope with the expected number of electrons in the apo- and holo- structures. If we disregard any cross-contamination during sample preparation (§3, ¶3), we would expect perfect clustering to result in four electron density maps where all ligands have occupancy of zero or unity. The observed occupancies were 0.18 & 0.00 (native), 0.62 & 0.03 (BEN+/NAG-), 0.09 & 1.00 (BEN-/NAG+) and 1.00 & 0.67 (BEN+/NAG+). The observed occupancies are, on average, 12.6% different from the ideal values of zero and unity.

#### A.2. Data Reduction and Structure Solving Statistics

The data reduction and structure solving statistics are given in Tables A1, A2 and A3. Because the largest clusters merge structures with significantly different ligands some of the R-factors degrade rather than improve. This is only to be expected. See (Kleywegt & Jones, 1997) for more information on R*_free_* and R*_work_*.

## Acknowledgements

Our thanks to Frances C. Bernstein for careful copy-editing and helpful suggestions. Work supported in part by

• US Dept. of Energy, Office of Science, DE-AC02-98CH10886, E-SC0012704, and KP1607011

• US NIH National Institute of General Medical Sciences, P41RR012408, P41GM103473, P41GM111244, and P30GM133893

• HJB supported in part by Dectris, Ltd.

---

**Fig. A1.**
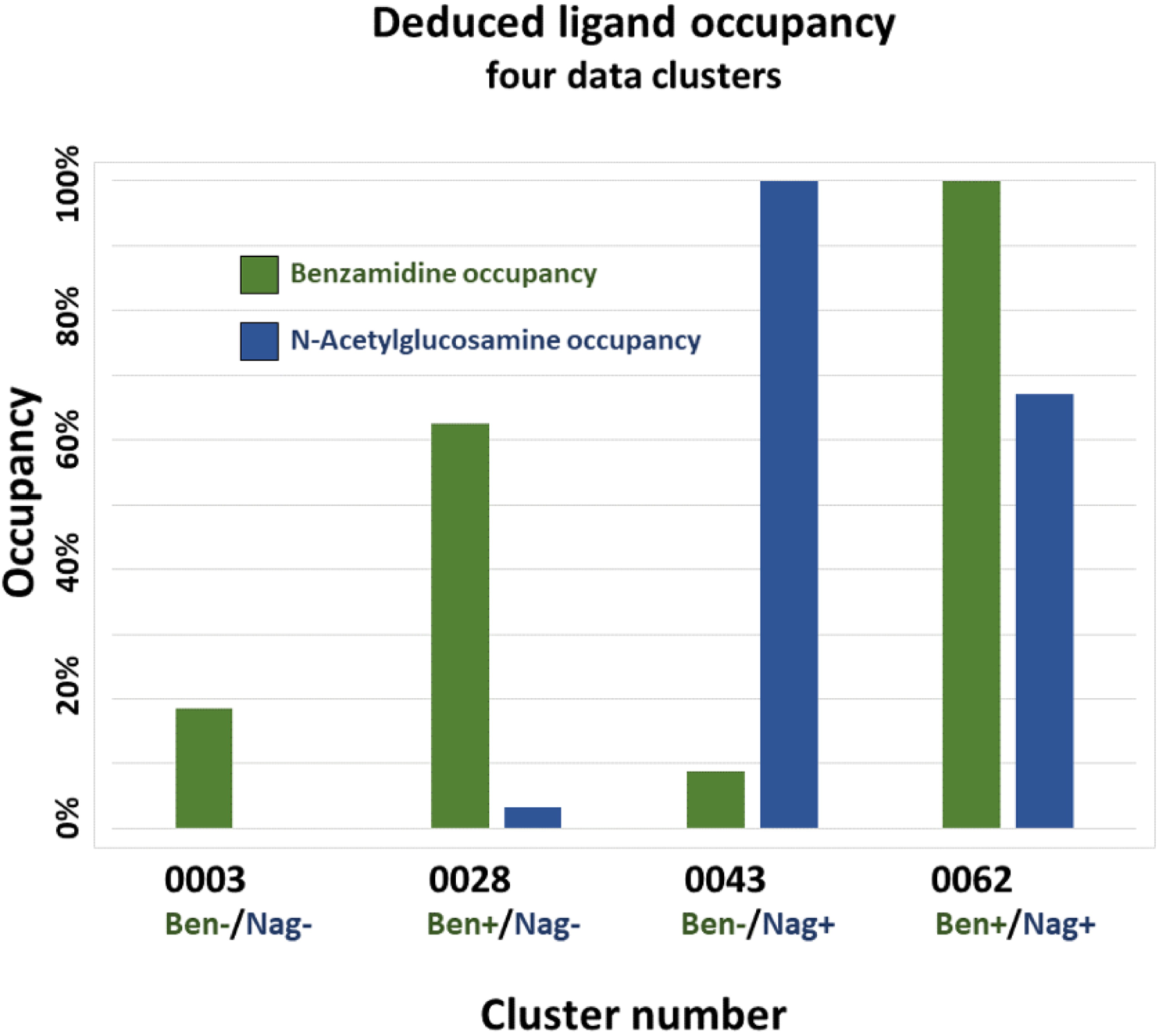
Occupancy of benzamidine (BEN) and N-Acetylglucosamine (NAG) ligands deduced from the integrated electron density in the volume occupied by each ligand. The data clusters were obtained using two-stage clustering (BLEND^NCDist^ followed by BLEND^SFDist^)

**Fig. A2.**
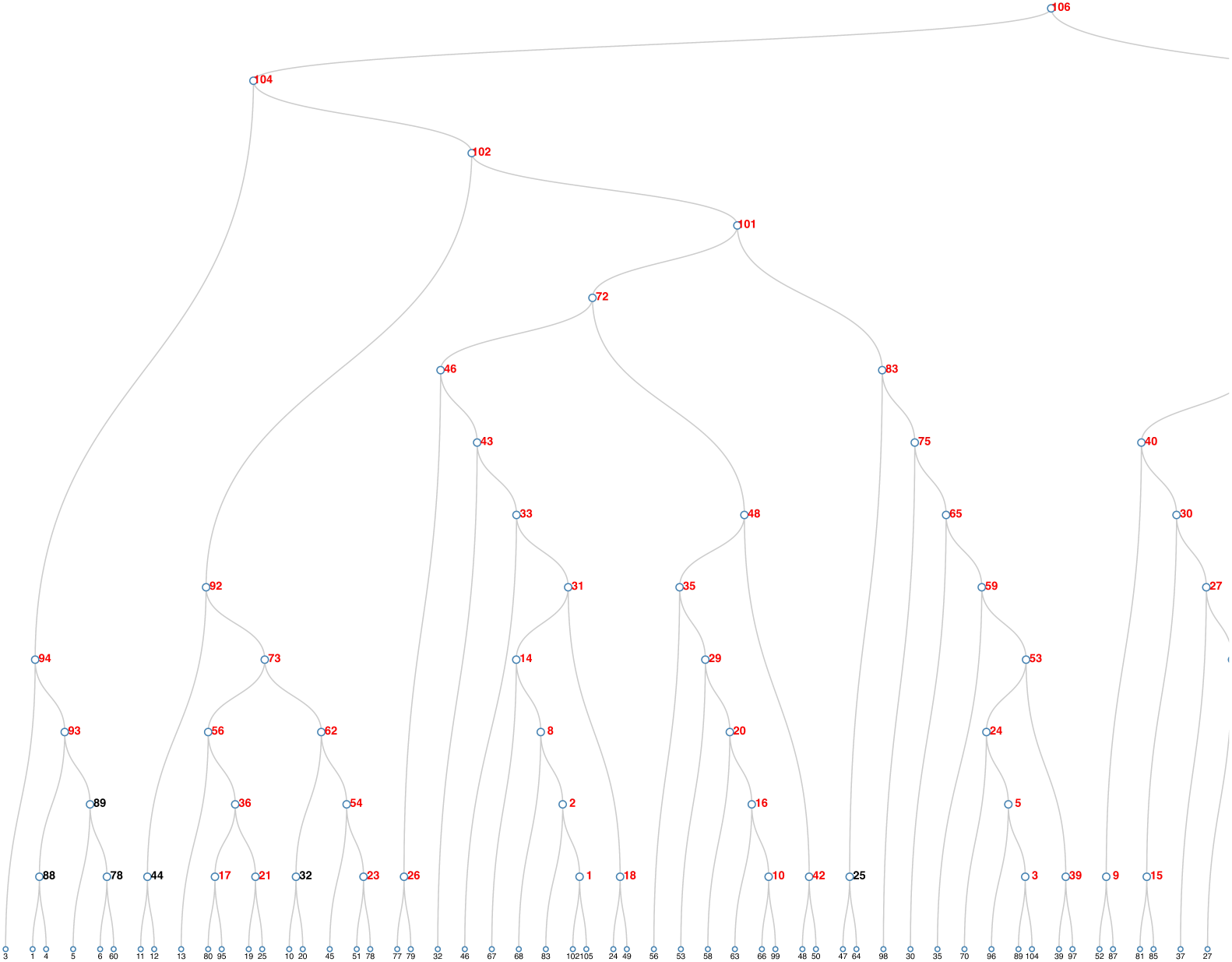
Left half of the complete dendrogram for all the data clusters that were obtained using two-stage clustering (BLEND^NCDist^ followed by BLEND^SFDist^). The other half of this dendrogram is shown in Fig. A3

**Fig. A3.**
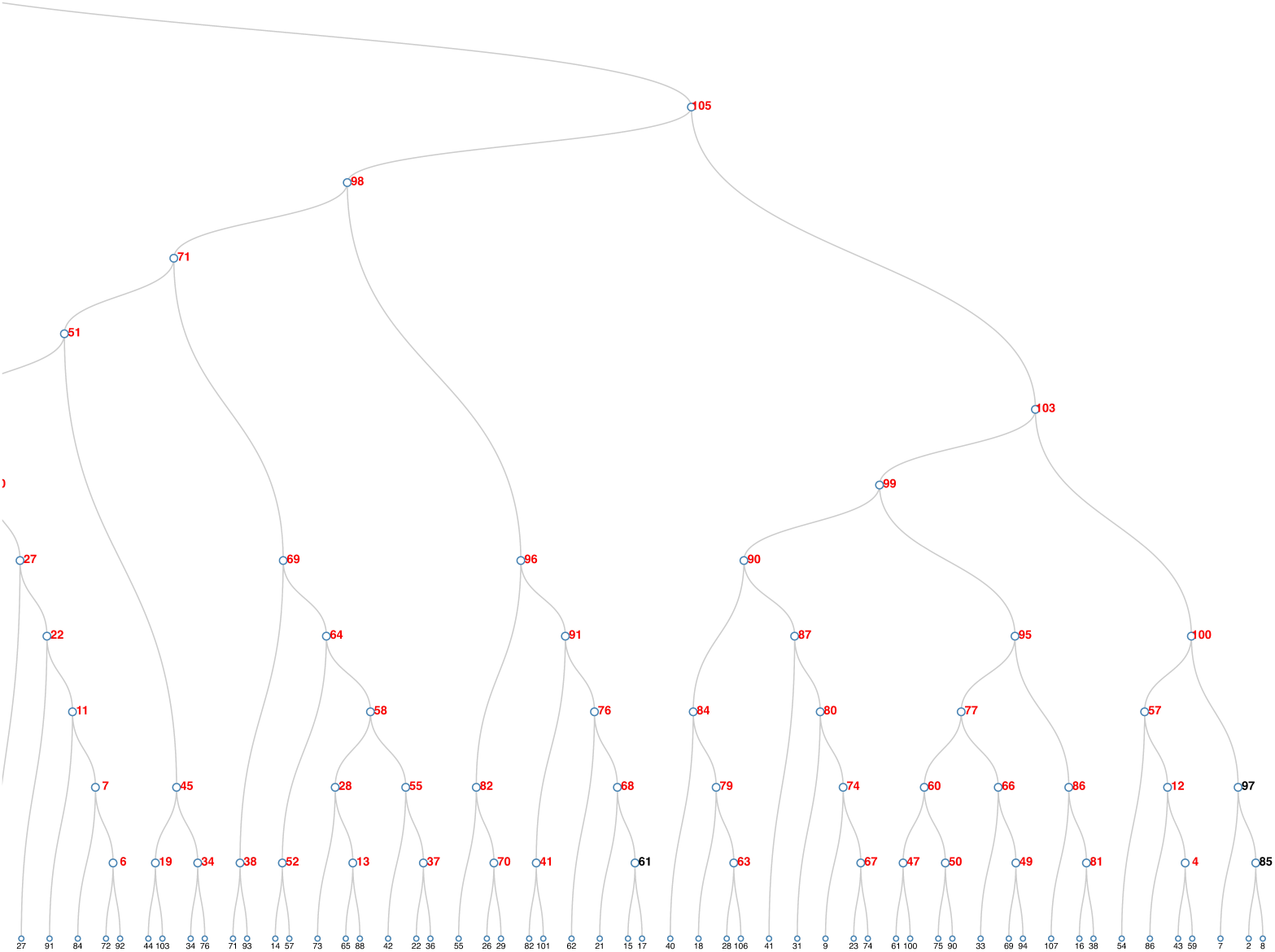
Right half of the complete dendrogram for all the data clusters that were obtained using two-stage clustering (BLEND^NCDist^ followed by BLEND^SFDist^). The other half of this dendrogram is shown in Fig. A2

**Fig. A4.**
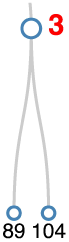
Portion of the dendrogram in Figs. A2 and A3 showing cluster 3

**Fig. A5.**
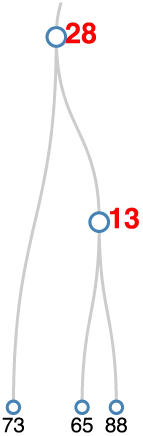
Portion of the dendrogram in Figs. A2 and A3 showing cluster 28

**Fig. A6.**
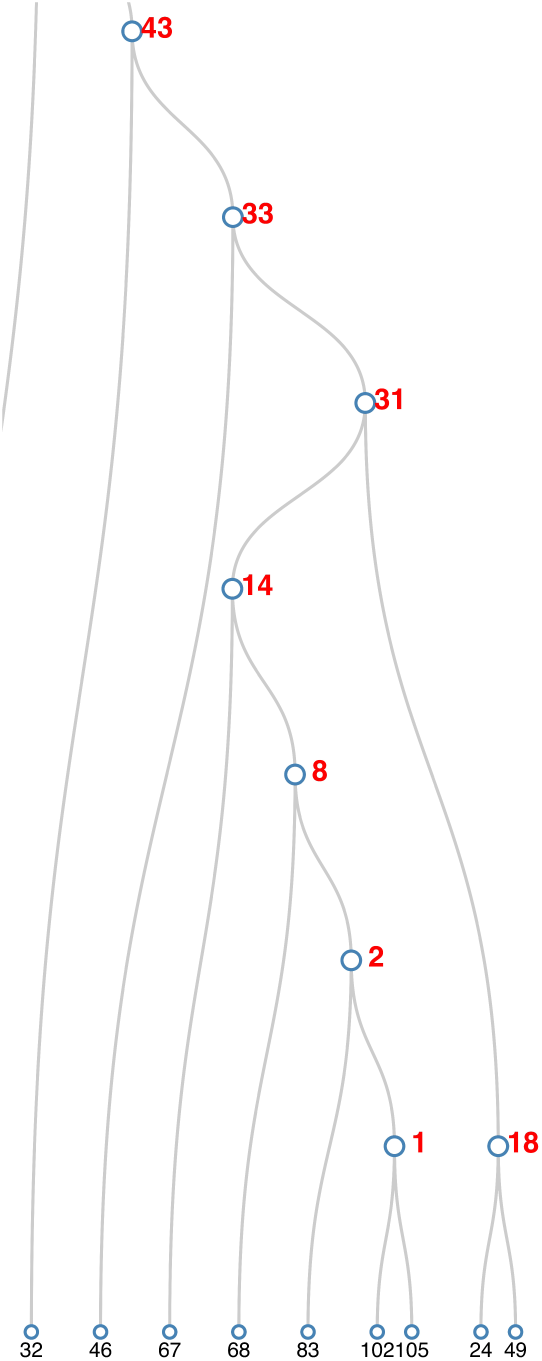
Portion of the dendrogram in Figs. A2 and A3 showing cluster 43

**Fig. A7.**
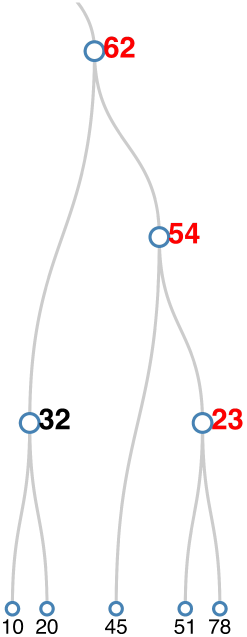
Portion of the dendrogram in Figs. A2 and A3 showing cluster 62

**Table A1.** Data reduction and structure solving statistics. Data are shown for all clusters determined using two factor clustering (NCDist followed by SFDist), continued in Tables A2 and A3

| Cluster<br>Number | Rmerge<br>(%) | Comp.<br>(%) | RefNum | R<sub>work</sub><br>Overall | R<sub>free</sub><br>Overall | R<sub>work</sub><br>Last shell | R<sub>free</sub><br>Last shell | CC<sub>work</sub><br>Overall | CC<sub>free</sub><br>Overall |
| --- | --- | --- | --- | --- | --- | --- | --- | --- | --- |
| 001 | 19.80 | 97.12 | 7740 | 0.172 | 0.230 | 0.204 | 0.266 | 0.950 | 0.916 |
| 002 | 22.60 | 99.19 | 7844 | 0.169 | 0.218 | 0.197 | 0.263 | 0.952 | 0.920 |
| 003 | 36.00 | 98.28 | 7909 | 0.212 | 0.286 | 0.345 | 0.349 | 0.946 | 0.904 |
| 004 | 19.30 | 92.99 | 7337 | 0.201 | 0.258 | 0.223 | 0.307 | 0.952 | 0.927 |
| 005 | 39.50 | 99.44 | 8004 | 0.208 | 0.282 | 0.331 | 0.336 | 0.945 | 0.899 |
| 006 | 28.50 | 96.13 | 7826 | 0.204 | 0.271 | 0.357 | 0.265 | 0.952 | 0.915 |
| 007 | 31.70 | 98.99 | 8059 | 0.196 | 0.253 | 0.327 | 0.345 | 0.953 | 0.930 |
| 008 | 23.00 | 99.62 | 7942 | 0.162 | 0.216 | 0.187 | 0.262 | 0.955 | 0.927 |
| 009 | 35.90 | 93.97 | 7580 | 0.235 | 0.315 | 0.368 | 0.333 | 0.935 | 0.887 |
| 010 | 28.70 | 98.60 | 7875 | 0.203 | 0.266 | 0.223 | 0.321 | 0.925 | 0.888 |
| 011 | 38.00 | 99.57 | 8108 | 0.192 | 0.249 | 0.324 | 0.321 | 0.953 | 0.930 |
| 012 | 33.80 | 98.10 | 7744 | 0.201 | 0.258 | 0.286 | 0.309 | 0.950 | 0.925 |
| 013 | 39.10 | 95.42 | 7794 | 0.223 | 0.286 | 0.349 | 0.405 | 0.929 | 0.891 |
| 014 | 25.40 | 99.83 | 7926 | 0.161 | 0.214 | 0.180 | 0.215 | 0.954 | 0.925 |
| 015 | 38.80 | 95.85 | 7851 | 0.220 | 0.287 | 0.307 | 0.308 | 0.937 | 0.902 |
| 016 | 34.70 | 99.36 | 7920 | 0.200 | 0.263 | 0.217 | 0.281 | 0.927 | 0.888 |
| 017 | 25.60 | 96.77 | 7770 | 0.179 | 0.235 | 0.236 | 0.299 | 0.944 | 0.921 |
| 018 | 30.20 | 90.16 | 7158 | 0.185 | 0.249 | 0.249 | 0.300 | 0.941 | 0.898 |
| 019 | 44.30 | 96.72 | 7800 | 0.234 | 0.296 | 0.332 | 0.384 | 0.936 | 0.896 |
| 020 | 32.40 | 99.85 | 7963 | 0.187 | 0.258 | 0.179 | 0.241 | 0.937 | 0.895 |
| 021 | 16.70 | 91.34 | 7298 | 0.173 | 0.247 | 0.275 | 0.309 | 0.952 | 0.914 |
| 022 | 38.20 | 99.74 | 8123 | 0.188 | 0.247 | 0.327 | 0.327 | 0.955 | 0.934 |
| 023 | 28.20 | 91.79 | 7371 | 0.187 | 0.247 | 0.220 | 0.311 | 0.936 | 0.909 |
| 024 | 41.80 | 99.72 | 8039 | 0.209 | 0.281 | 0.332 | 0.348 | 0.942 | 0.894 |
| 026 | 21.50 | 91.55 | 7286 | 0.173 | 0.240 | 0.212 | 0.246 | 0.946 | 0.899 |
| 027 | 39.80 | 99.83 | 8130 | 0.186 | 0.243 | 0.312 | 0.297 | 0.955 | 0.936 |
| 028 | 42.60 | 97.78 | 7988 | 0.212 | 0.271 | 0.352 | 0.403 | 0.940 | 0.915 |
| 029 | 36.10 | 99.85 | 7983 | 0.189 | 0.257 | 0.180 | 0.259 | 0.934 | 0.889 |
| 030 | 43.10 | 99.94 | 8140 | 0.178 | 0.233 | 0.264 | 0.282 | 0.956 | 0.938 |
| 031 | 27.90 | 99.89 | 7933 | 0.159 | 0.212 | 0.177 | 0.216 | 0.954 | 0.926 |
| 033 | 27.10 | 99.95 | 7938 | 0.157 | 0.209 | 0.164 | 0.214 | 0.954 | 0.928 |
| 034 | 54.40 | 94.52 | 7685 | 0.235 | 0.297 | 0.376 | 0.396 | 0.930 | 0.885 |
| 035 | 38.50 | 99.87 | 7985 | 0.196 | 0.266 | 0.190 | 0.251 | 0.929 | 0.880 |
| 036 | 24.40 | 99.25 | 7949 | 0.167 | 0.212 | 0.211 | 0.257 | 0.950 | 0.935 |
| 037 | 45.50 | 83.97 | 6854 | 0.223 | 0.311 | 0.331 | 0.505 | 0.946 | 0.904 |
| 038 | 48.50 | 96.99 | 7990 | 0.260 | 0.316 | 0.402 | 0.386 | 0.919 | 0.890 |
| 039 | 44.20 | 96.51 | 7770 | 0.231 | 0.306 | 0.353 | 0.291 | 0.930 | 0.879 |
| 040 | 43.40 | 99.97 | 8141 | 0.181 | 0.228 | 0.267 | 0.317 | 0.954 | 0.942 |
| 041 | 60.10 | 96.72 | 7938 | 0.259 | 0.345 | 0.362 | 0.421 | 0.901 | 0.829 |
| 042 | 27.60 | 91.75 | 7315 | 0.200 | 0.271 | 0.268 | 0.325 | 0.932 | 0.884 |
| 043 | 28.10 | 99.98 | 7940 | 0.159 | 0.210 | 0.161 | 0.212 | 0.953 | 0.925 |
| 045 | 56.40 | 99.58 | 8097 | 0.214 | 0.270 | 0.353 | 0.358 | 0.944 | 0.916 |
| 046 | 28.40 | 99.98 | 7940 | 0.158 | 0.210 | 0.167 | 0.225 | 0.954 | 0.927 |
| 047 | 51.10 | 97.24 | 7842 | 0.278 | 0.362 | 0.390 | 0.370 | 0.917 | 0.851 |

**Table A2.** Data reduction and structure solving statistics. Data are shown for all clusters determined using two factor clustering (NCDist followed by SFDist), continued from Table

| Cluster<br>Number | Rmerge<br>(%) | Comp.<br>(%) | RefNum | R<sub>work</sub><br>Overall | R<sub>free</sub> | R<sub>work</sub><br>Last shell | R<sub>free</sub> | CC<sub>work</sub><br>Overall | CC<sub>free</sub> |
| --- | --- | --- | --- | --- | --- | --- | --- | --- | --- |
| 048 | 40.70 | 99.98 | 7972 | 0.193 | 0.267 | 0.183 | 0.258 | 0.930 | 0.883 |
| 049 | 46.50 | 96.21 | 7771 | 0.272 | 0.350 | 0.416 | 0.404 | 0.910 | 0.842 |
| 050 | 52.70 | 96.94 | 7834 | 0.273 | 0.350 | 0.359 | 0.432 | 0.908 | 0.848 |
| 051 | 48.60 | 99.97 | 8141 | 0.177 | 0.230 | 0.253 | 0.309 | 0.955 | 0.939 |
| 052 | 40.30 | 92.89 | 7586 | 0.233 | 0.310 | 0.381 | 0.407 | 0.932 | 0.876 |
| 053 | 49.60 | 99.92 | 8053 | 0.204 | 0.275 | 0.297 | 0.296 | 0.944 | 0.898 |
| 054 | 39.80 | 97.44 | 7861 | 0.212 | 0.278 | 0.269 | 0.395 | 0.918 | 0.882 |
| 055 | 49.40 | 95.61 | 7812 | 0.222 | 0.304 | 0.343 | 0.446 | 0.945 | 0.909 |
| 056 | 24.90 | 99.50 | 7940 | 0.163 | 0.214 | 0.197 | 0.267 | 0.951 | 0.933 |
| 057 | 38.40 | 98.92 | 7810 | 0.204 | 0.261 | 0.273 | 0.337 | 0.948 | 0.923 |
| 058 | 54.80 | 99.63 | 8137 | 0.206 | 0.265 | 0.341 | 0.400 | 0.942 | 0.919 |
| 059 | 52.10 | 99.98 | 8052 | 0.202 | 0.276 | 0.282 | 0.296 | 0.942 | 0.895 |
| 060 | 55.80 | 99.57 | 8049 | 0.261 | 0.349 | 0.387 | 0.426 | 0.925 | 0.860 |
| 062 | 40.80 | 99.28 | 7919 | 0.193 | 0.264 | 0.230 | 0.340 | 0.935 | 0.901 |
| 063 | 48.70 | 96.83 | 7875 | 0.314 | 0.408 | 0.348 | 0.479 | 0.877 | 0.789 |
| 064 | 57.90 | 99.91 | 8165 | 0.202 | 0.259 | 0.333 | 0.370 | 0.941 | 0.920 |
| 065 | 53.20 | 99.99 | 8039 | 0.204 | 0.276 | 0.288 | 0.334 | 0.942 | 0.898 |
| 066 | 50.70 | 98.52 | 7985 | 0.270 | 0.350 | 0.404 | 0.366 | 0.914 | 0.849 |
| 067 | 49.90 | 96.12 | 7726 | 0.299 | 0.364 | 0.290 | 0.388 | 0.888 | 0.853 |
| 068 | 59.40 | 92.95 | 7526 | 0.257 | 0.338 | 0.388 | 0.384 | 0.914 | 0.857 |
| 069 | 61.50 | 99.92 | 8166 | 0.201 | 0.259 | 0.324 | 0.371 | 0.938 | 0.918 |
| 070 | 61.50 | 87.62 | 7190 | 0.291 | 0.363 | 0.403 | 0.233 | 0.897 | 0.833 |
| 071 | 54.10 | 99.92 | 8166 | 0.175 | 0.228 | 0.233 | 0.247 | 0.954 | 0.937 |
| 072 | 43.10 | 99.99 | 7941 | 0.181 | 0.234 | 0.185 | 0.285 | 0.941 | 0.905 |
| 073 | 36.10 | 99.99 | 7976 | 0.168 | 0.234 | 0.194 | 0.283 | 0.946 | 0.918 |
| 074 | 62.40 | 97.86 | 7855 | 0.289 | 0.363 | 0.315 | 0.404 | 0.893 | 0.853 |
| 075 | 55.90 | 99.99 | 8039 | 0.204 | 0.276 | 0.284 | 0.336 | 0.940 | 0.893 |
| 076 | 63.00 | 97.94 | 7931 | 0.261 | 0.340 | 0.376 | 0.328 | 0.914 | 0.854 |
| 077 | 67.00 | 99.98 | 8107 | 0.280 | 0.335 | 0.381 | 0.338 | 0.904 | 0.872 |
| 079 | 89.50 | 99.00 | 7974 | 0.377 | 0.417 | 0.440 | 0.501 | 0.740 | 0.710 |
| 080 | 64.40 | 98.94 | 7941 | 0.291 | 0.362 | 0.325 | 0.384 | 0.896 | 0.847 |
| 082 | 59.70 | 98.13 | 8053 | 0.281 | 0.369 | 0.363 | 0.424 | 0.899 | 0.845 |
| 083 | 56.60 | 99.99 | 8039 | 0.206 | 0.276 | 0.267 | 0.340 | 0.936 | 0.891 |
| 084 | 78.80 | 99.49 | 8014 | 0.372 | 0.411 | 0.395 | 0.424 | 0.790 | 0.760 |
| 086 | 64.20 | 98.45 | 7904 | 0.313 | 0.372 | 0.415 | 0.452 | 0.868 | 0.810 |
| 087 | 69.30 | 99.52 | 7988 | 0.290 | 0.364 | 0.328 | 0.392 | 0.896 | 0.847 |
| 090 | 90.30 | 99.89 | 8020 | 0.358 | 0.408 | 0.367 | 0.472 | 0.791 | 0.737 |
| 091 | 71.00 | 99.75 | 8079 | 0.249 | 0.341 | 0.353 | 0.378 | 0.911 | 0.846 |
| 092 | 35.20 | 99.99 | 7976 | 0.165 | 0.229 | 0.188 | 0.241 | 0.948 | 0.919 |
| 093 | 36.20 | 90.95 | 7230 | 0.225 | 0.296 | 0.334 | 0.341 | 0.923 | 0.863 |
| 094 | 36.50 | 92.16 | 7324 | 0.221 | 0.292 | 0.319 | 0.338 | 0.923 | 0.864 |
| 095 | 64.00 | 99.95 | 8105 | 0.267 | 0.354 | 0.392 | 0.355 | 0.916 | 0.855 |
| 096 | 74.70 | 99.91 | 8092 | 0.243 | 0.329 | 0.349 | 0.386 | 0.915 | 0.853 |

**Table A3.**
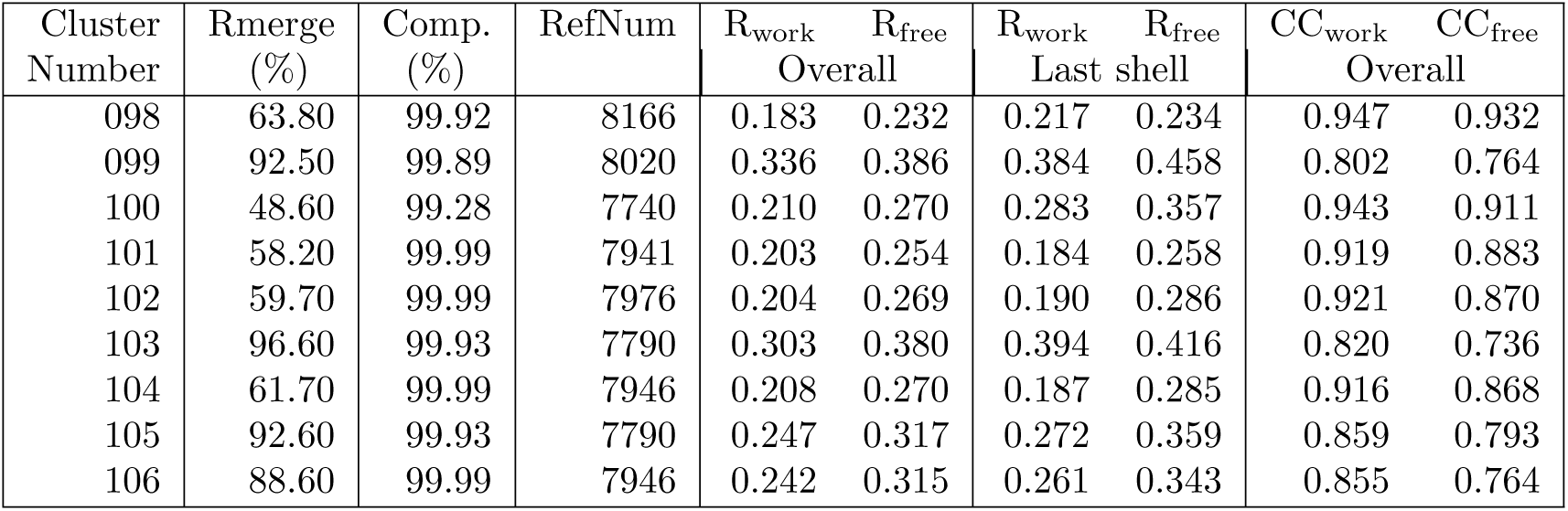
Data reduction and structure solving statistics. Data are shown for all clusters determined using two factor clustering (NCDist followed by SFDist), continued from Tables

## Synopsis

The effectiveness of clustering to merge datasets from large numbers of crystals in serial crystallography can be improved by combining multiple clustering techniques using cell parameter-based clustering for very incomplete sets and switching to reflection-based clustering once preliminary merging has increased completeness.

